# Impaired S-nitrosylation of Cx43 prevents arrhythmogenicity and myocardial injury upon cardiac stress in Duchenne Muscular Dystrophy

**DOI:** 10.1101/2024.08.29.610357

**Authors:** Manuel F. Muñoz, Jonathan J. Quan, Thao T. Nguyen, Janet Nuno, Adrian Sheehy, Pia C. Burboa, Pablo S. Gaete, Mauricio A. Lillo, Jorge E. Contreras

## Abstract

Connexin-43 (Cx43) plays a critical role in the propagation of action potentials and cardiac contractility. In healthy cardiomyocytes, Cx43 is mainly located at the intercalated disk; however, Cx43 remodeling is observed in cardiac pathologies and is linked with arrhythmogenesis and sudden cardiac death. Using a mouse model of Duchenne muscular dystrophy (DMD), we previously demonstrated that Cx43 localizes to the lateral side of dystrophic cardiomyocytes, forming undocked hemichannels. β-adrenergic signaling-induced cardiac stress promotes S-nitrosylation and the opening of undocked Cx43 hemichannels leading to disrupted cardiac membrane excitability and deadly arrhythmogenic behaviors. To establish the direct role of S-nitrosylated Cx43 in DMD cardiomyopathy, we generated knockin DMD^mdx^ mice with reduced levels of S-nitrosylated Cx43, by replacing cysteine 271 with a serine in one Cx43 of the unique site for S-nitrosylation of Cx43 (DMD^mdx^:C271S^+/-^). Immunofluorescence analysis revealed that cardiac Cx43 lateralization in DMD^mdx^:C271S^+/-^ mice was similar to DMD^mdx^ mice, indicating that the genetic modification did not prevent Cx43 remodeling. Upon isoproterenol treatment, DMD^mdx^ mice displayed a higher incidence of arrhythmogenic events when compared to DMD^mdx^:C271S^+/-^ mice, which more closely resemble wild-type mice. Optical mapping imaging in isolated hearts showed that DMD^mdx^ mice displayed aberrant Ca^2+^ signaling and prolonged action potentials, which is restored in DMD^mdx^:C271S^+/-^ mice. Isoproterenol treatment evoked severe myocardial injury in DMD^mdx^ mice, which was significantly attenuated in DMD^mdx^:C271S^+/-^ mice. Notably, DMD^mdx^ mice treated with Gap19, a Cx43 hemichannel blocker, exhibited cardioprotection against myocardial injury. We concluded that S-nitrosylation of Cx43 proteins is a fundamental NO-mediated mechanism involved in arrhythmias and myocardial injury in DMD^mdx^, occurring through the opening of hemichannels following β-adrenergic stress.

## INTRODUCTION

Duchenne muscular dystrophy (DMD) is a severe and progressive muscle-wasting disease that primarily affects males, occurring in approximately 1 in 5,000 male newborns (1, 2). It is characterized by an X-linked mutation leading to the loss of function in the dystrophin gene, which encodes a crucial protein responsible for maintaining membrane stability in muscle cells (2). DMD patients typically present with muscle weakness, respiratory failure, and cardiac dysfunction, with cardiomyopathies, particularly dilated cardiomyopathy (3), being the leading cause of mortality (1). Among the primary clinical heart complications observed in these patients are acute myocardial injury (4, 5), elevated plasma levels of cardiac troponin-I (cTnI) (6), and significant sympathetic activity (7). Despite extensive research, the molecular mechanisms underlying DMD cardiomyopathy remain incompletely understood. Current therapeutic approaches, such as gene therapy using adeno-associated virus (AAV)-based methods to restore dystrophin (8), losartan treatment to antagonize angiotensin II type I receptors (9–11), and β-receptor blocker therapy (12) have only partially reduced the cardiac dysfunction and associated mortality in these patients. This represents a critical unsolved challenge that must be addressed to improve the lives of DMD patients.

In the heart, connexin-43 (Cx43) is the predominant connexin isoform expressed in the ventricular myocardium (13, 14). Intriguingly, Cx43 is extensively remodeled in DMD cardiomyopathy, exhibiting abnormal lateral distribution in cardiomyocytes (15–19). Our laboratory has demonstrated that hyperactivity of remodeled Cx43 hemichannels critically contributes to arrhythmogenicity in a dystrophic mouse model (DMD^mdx^) (18–20). We reported that DMD^mdx^ mice with lower levels of Cx43 proteins (DMD^mdx^ mice: Cx43^+/-^) (20) or the blockade of Cx43 hemichannels using connexin mimetic peptides displayed a low incidence of arrhythmias and sudden cardiac death following isoproterenol challenge (21). Cx43 hemichannel blockade not only prevented cardiac stress-elicited triggered activity (TA) in DMD^mdx^ cardiomyocytes (19) but also demonstrated improvements in infarct size and cardiomyocyte viability following ischemia/reperfusion *in vivo* and *in vitro* (22). This confirms that Cx43 hemichannel hyperactivity is pivotal in mediating these pro-arrhythmic events and affecting myocardial cell viability.

Our laboratory and another group reported that stimulation with isoproterenol promotes hyper-nitrosylation of Cx43 proteins in the hearts of DMD^mdx^ mice (19, 23), which were prevented by treating the animals with the nitric oxide (NO) synthase inhibitor, N(ω)-nitro-L-arginine methyl ester (L-NAME) (19). L-NAME also prevented isoproterenol-induced arrhythmias and TAs in DMD^mdx^ mice. However, this treatment equally affects S-nitrosylation levels of other key proteins implicated in sarcoplasmic reticulum Ca^2+^ leaks and ventricular arrhythmias including ryanodine receptor type 2 (RyR2) (24), Ca^2+^/calmodulin-dependent protein kinase II (CaMKII) (25), voltage-gated Na^+^ channels (26), and other proteins associated with cardiac dysfunction and Ca^2+^ handling (27). Thus, the precise role of Cx43 S-nitrosylation in DMD-related cardiomyopathy remains unresolved. Electrophysiological studies using *in vitro* heterologous expression systems revealed that NO induces S-nitrosylation of Cx43 at cysteine 271 (C271), which is critical for promoting the opening of Cx43 hemichannel (19). Therefore, we hypothesize that S-nitrosylation of lateralized Cx43 hemichannels at C271 is critical for promoting excessive Ca^2+^ influx, membrane depolarization, and triggered activity after β-adrenergic stimulation in DMD^mdx^ cardiomyocytes. To test this hypothesis, we developed a novel DMD^mdx^ knock-in mouse model with reduced levels of S-nitrosylated Cx43 (DMD^mdx^:C271S^+/-^). Our results showed that the specific impairing S-nitrosylation of Cx43 in DMD^mdx^ mice is sufficient to restore cardiac excitability and prevent lethal arrhythmias and myocardial injury.

## RESULTS

### Generation and characterization of C271S^+/-^ and DMD^mdx^: C271S^+/-^ mouse lines

We have previously shown that NO-induced Cx43 hemichannel activation via S-nitrosylation occurs at the residue C271 (19). To evaluate how the substitution of C271 with serine affects Cx43 hemichannel activation, we measured Cx43 wild-type and mutant (C271S) hemichannel ionic currents in Xenopus oocytes using the two-electrode voltage clamp technique (28). Wild-type Cx43-expressing oocytes displayed increased ionic currents when exposed to the NO donor 100 µM DEA NONOate at potentials close to cardiomyocyte membrane potentials (-70 mV) (18) (Supplementary Figure 1). In contrast, oocytes expressing either the Cx43 mutant (Cx43^C271S^) or co-expressing both the wild-type and C271S mutant (Cx43^WT/C271S^) displayed no significant ionic currents under the same conditions. This indicates that the C271S mutation abolishes NO-induced activation of Cx43 hemichannels.

To assess the *in vivo* effects of S-nitrosylated Cx43 hemichannels at cysteine 271, we developed a mouse model in which this site was replaced with serine (C271S) using CRISPR-Cas9 genome editing (Supplementary Figure 2A). The generation of the C271S mutation produced wild-type (C271S^-/-^), heterozygotes (C271S^+/-^), and homozygotes C271S (C271S^+/+^) mice. Supplementary Figure 2B shows a representative biotin switch assay that confirms reduced or zero levels of S-nitrosylation in C271S^+/-^ and C271S^+/+^ mice, respectively. Importantly, C271S^+/-^ and C271S^+/+^ mice showed no difference in animal weight, heart weight, and heart/animal weight ratio with respect to wild-type mice (Supplementary Figure 2C, 2D, and 2E). However, C271S^+/+^ mice (138.5 ± 3.14 mmHg) displayed hypertension in comparison to wild-type (101.5 ± 0.92 mmHg) and C271S^+/-^ mice (105.6 ± 3.74 mmHg), indicating off-target effects that might eventually affect cardiac function. This precludes the use of C271S^+/+^ mice for rigorously establishing the relationship between S-nitrosylated Cx43 hemichannels and DMD^mdx^ cardiomyopathy. Based on our electrophysiological findings the C271S mutation serves as a dominant negative for NO-dependent Cx43 hemichannel opening (Supplementary Figure 1), we generated a DMD^mdx^ knockin mouse with reduced S-nitrosylation of Cx43 using the heterozygote C271S^+/-^ mice. This model is referred to as DMD^mdx^: C271S^+/-^ mice (Supplementary Figure 3). Importantly, we observed no weight differences among 4-6-month-old wild-type, DMD^mdx^, and DMD^mdx^: C271S^+/-^ mice (Supplementary Figure 4). Remarkably, DMD^mdx^: C271S^+/-^ mice did not develop the hypertrophy characteristic of DMD^mdx^, exhibiting heart weight and heart-to-body weight ratios comparable to those of wild-type and C271S^+/-^ mouse lines (Supplementary Figure 4) suggesting a protective effect against hypertrophy.

### β-adrenergic stimulation evokes severe arrhythmias in DMD^mdx^ mice, which are prevented in DMD^mdx^:C271S^+/-^ mice

Arrhythmogenicity observed in DMD^mdx^ hearts upon β-adrenergic stimulation correlates with increased S-nitrosylation of Cx43, both of which are inhibited by L-NAME, a NOS blocker (19). However, the specific impact of reduced S-nitrosylation of Cx43 at cysteine 271 on arrhythmogenesis during isoproterenol-induced cardiac stress *in vivo* has not been explored. To investigate this, we performed electrocardiogram (ECG) telemetry recordings to analyze arrhythmogenic events in wild-type, DMD^mdx^, and DMD^mdx^:C271S^+/-^ mice under control conditions and for 24 hours post-isoproterenol injection (5 mg/kg). Representative ECG recordings at baseline, 1 hour, and 24 hours post-isoproterenol from wild-type, DMD^mdx^, and DMD^mdx^:C271S^+/-^ mice are shown in Figure 1A. Figure 1B shows the increased number of total arrhythmogenic events during the 24 hours post-isoproterenol in DMD^mdx^ mice (212 ± 42.21 events). In contrast, DMD^mdx^: C271S^+/-^ mice (92.4 ± 18.38 events) displayed a significantly lower number of arrhythmogenic events, comparable to the number observed in wild-type mice (31.14 ± 5.81 events). The total arrhythmogenic events included single premature ventricular contractions (PVCs) (Figure 1C, left panel), double PVCs (Figure 1C, middle panel), and non-sustained ventricular tachycardia (NSVT) (Figure 1C, right panel). Collectively, our data demonstrate that DMD^mdx^: C271S^+/-^ mice exhibit anti-arrhythmic protection against β-adrenergic-mediated cardiac stress observed in DMD^mdx^ mice.

**Figure 1.**
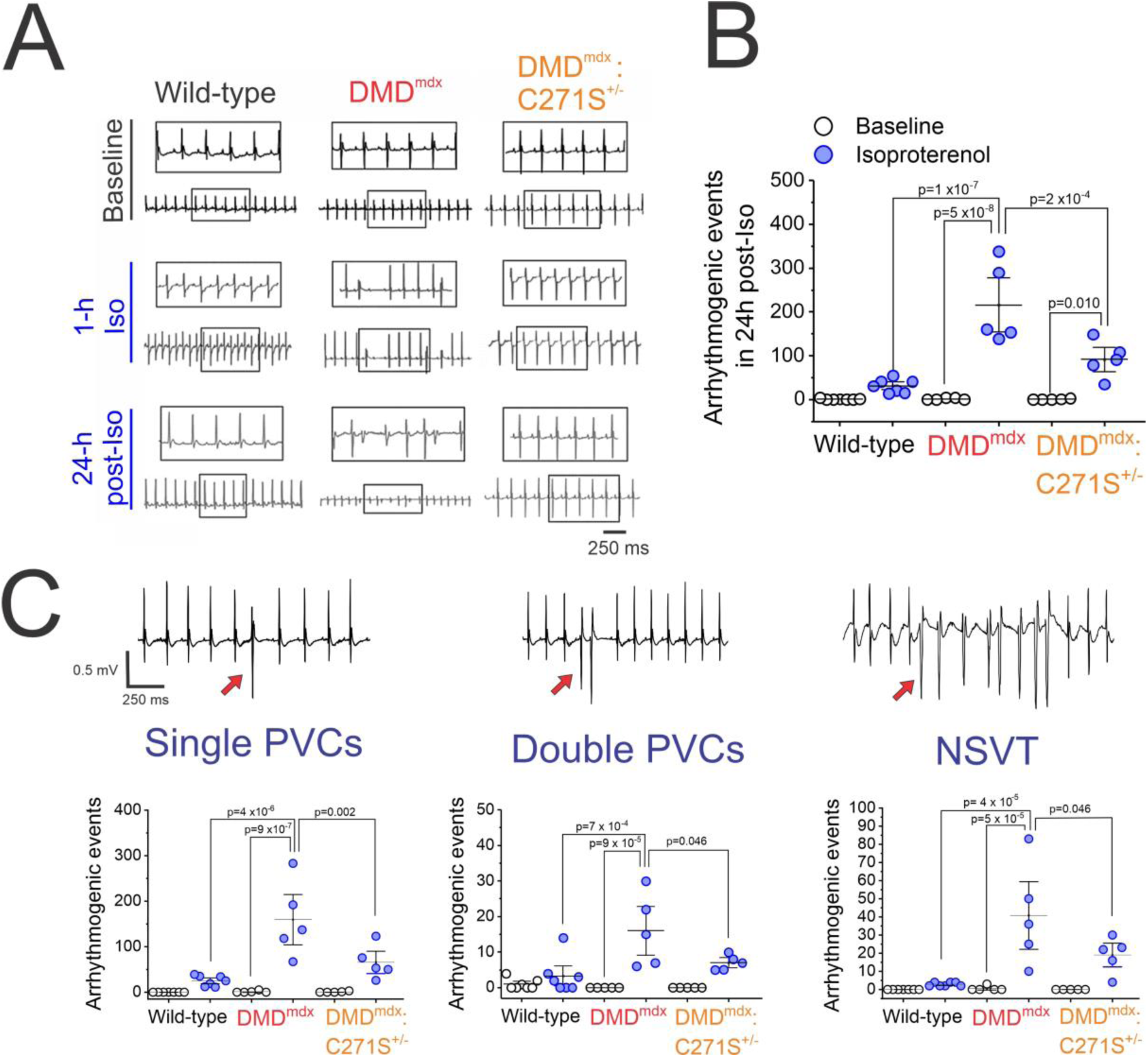
Reduced levels of S-nitrosylation of Cx43 hemichannels rescue the deadly DMD arrhythmogenicity induced by β-adrenergic stimulation-mediated cardiac stress. (A) Representative electrocardiogram (ECG) recordings from wild-type, DMD^mdx^, and DMD^mdx^: C271S^+/-^ mice via telemetry implants. The top trace represents recordings obtained at baseline. Traces in the middle and bottom indicate 1-hour and 24-h post-isoproterenol (5 mg/kg) challenge injected intraperitoneally, respectively. Scale bar = 250 ms. (B) Quantification of the total number of arrhythmogenic events before (baseline, white circles) and during the 24-hour post-isoproterenol injection (isoproterenol, blue circles). (C) Quantification of the type of arrhythmogenic events induced by β-adrenergic stimulation: single premature ventricular contractions (PVCs) (left), double PVCs (middle), and non-sustained ventricular tachycardia (NSVT) (three or more consecutive PVCs) (right). The type of arrhythmia can be graphically observed at the top of each graph, indicated by a red arrow. *n*=5-7 mice per group. Each dot represents an independent animal. Group comparisons were made using a one-way ANOVA followed by Tukey’s post hoc test. Data are presented as means ± SEM.

### Remodeling of Cx43 is not prevented in hearts from DMD^mdx^: C271S^+/-^ mice

We have previously demonstrated that Cx43 remodeling is mediated by hypophosphorylation of serine residues 325, 328, and 330 and that it is a factor critical for the development of cardiac dysfunction and arrhythmogenicity in DMD^mdx^ mice (18). Therefore, we investigated whether S-nitrosylation of Cx43 at residue C271 also contributes to this remodeling in DMD^mdx^ mice. To evaluate Cx43 distribution in the ventricles, we performed immunostaining for Cx43 (green), the intercalated disk marker N-cadherin (red), and the cell membrane marker wheat germ agglutinin (WGA) (blue) in ventricular sections of 4-6-month-old wild-type, DMD^mdx^, and DMD^mdx^: C271S^+/-^, and C271S^+/-^ mice. Representative confocal images (Figure 2A) from wild-type, DMD^mdx^, DMD^mdx^: C271S^+/-^, and C271S^+/-^ mice show that Cx43 is predominantly localized to the lateral side of cardiomyocytes in DMD^mdx^ and DMD^mdx^:C271S^+/-^ hearts, whereas it remains primarily at the intercalated disks in wild-type and C271S^+/-^ hearts. Quantification of Cx43 remodeling (Figure 2B) reveals a significantly higher percentage in DMD^mdx^ hearts compared to wild-type hearts, with mean remodeling percentages of 37.35 ± 1.70 % and 19.07 ± 3.56 %, respectively. C271S^+/-^ hearts exhibited values similar (22.77 ± 2.00 %) to wild-type, whereas DMD^mdx^:C271S^+/-^ hearts displayed remodeling percentages comparable to DMD^mdx^ hearts (38.99 ± 2.94 %). These findings indicate that lack of S-nitrosylation at residue C271 does not prevent Cx43 lateralization in DMD^mdx^ hearts and, suggesting that this is not the mechanism by which arrhythmias are ameliorate in DMD^mdx^:C271S^+/-^ mice.

**Figure 2.**
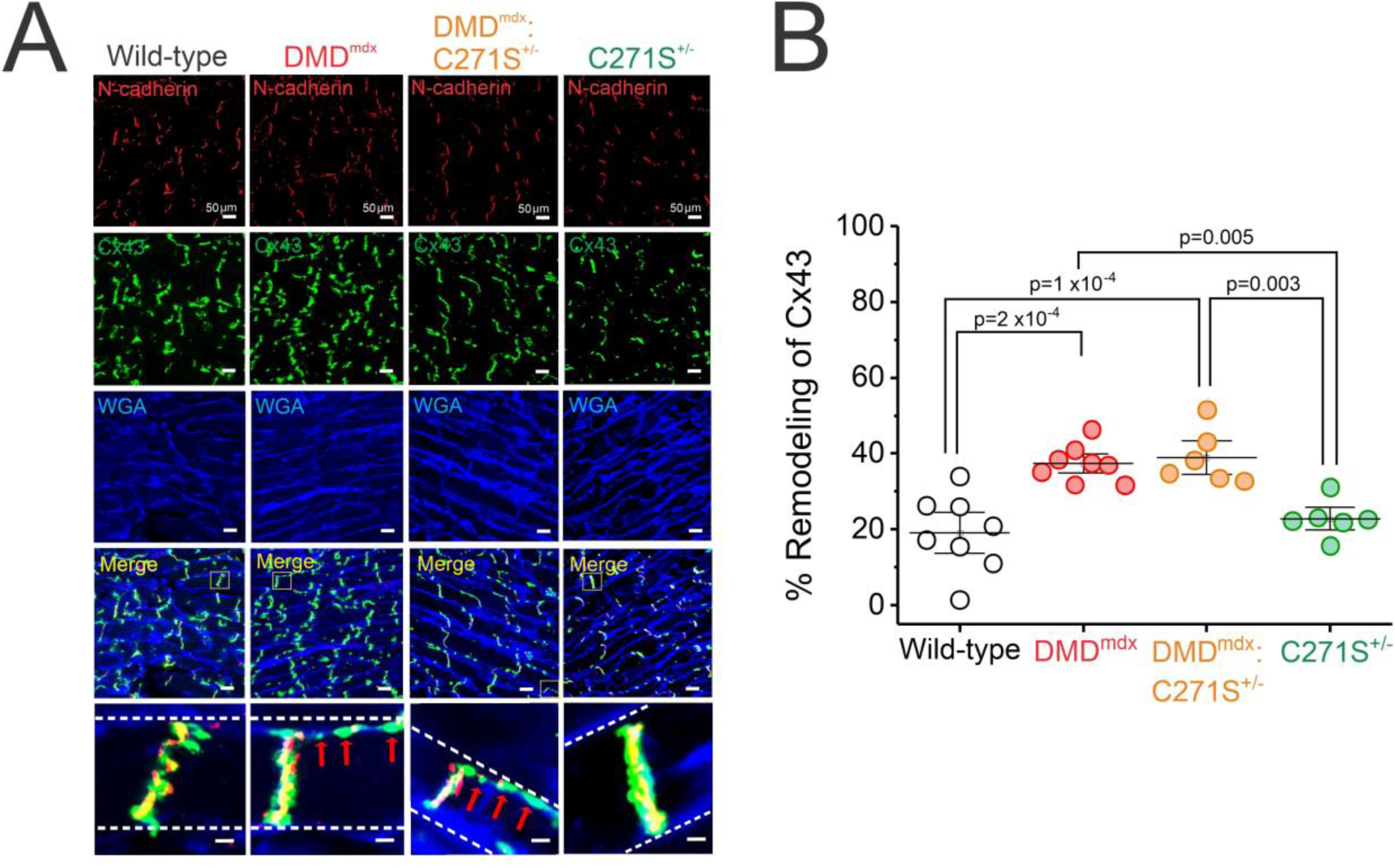
Remodeling of cardiac Cx43 is not prevented by decreased levels of S-nitrosylated Cx43 in DMD^mdx^ hearts (DMD^mdx^:C271S^+/-^). (A) Representative confocal images of immunofluorescence against N-cadherin (red) and Cx43 proteins (green) in wild-type, DMD^mdx^, DMD^mdx^:C271S^+/-^, and C271S^+/-^ mouse hearts. Plasma membranes were stained with wheat germ agglutinin (WGA) (blue). Scale bar = 50 μm. The bottom row shows inset areas (yellow squares) from merged photos of Cx43, N-cadherin (a marker of the intercalated disc), and WGA. Red arrows indicate positive signals for Cx43 at the lateral side of cardiomyocytes (DMD^mdx^ and DMD^mdx^:C271S^+/-^ hearts) (B) Percentage of remodeling of Cx43 in wild-type, DMD^mdx^, DMD^mdx^:C271S^+/-^, and C271S^+/-^ mouse heart sections, represented as the percentage of total mislocalized Cx43 with N-cadherin. *n*=6-8 mice per group. Each dot represents an independent animal. Group comparisons were made using a one-way ANOVA followed by Tukey’s post hoc test. Data are presented as means ± SEM.

### Prolonged action potential and aberrant Ca^2+^ signaling in DMD^mdx^ hearts are prevented in isolated cardiac hearts from DMD^mdx^:C271S^+/-^ mice

We previously demonstrated that β-adrenergic-induced opening of lateralized Cx43 hemichannels mediates altering cardiac excitability in DMD^mdx^ (19) and S3A cardiomyocytes (21). However, the specific role of Cx43 S-nitrosylation in this phenomenon remains unexplored. To investigate this, we performed optical mapping recordings to assess membrane potential changes (RH237) in Langendorff-perfused hearts at baseline and under isoproterenol (4 µM) stimulation during sinus rhythm in the left ventricles in wild-type, DMD^mdx^, and DMD^mdx^:C271S^+/-^ mice. Figures 3A and 3B show representative heat maps and traces of action potential durations (APDs) at 90% repolarization (APD_90_) for wild-type, DMD^mdx^, and DMD^mdx^:C271S^+/-^ hearts before (baseline) and after isoproterenol treatment. APDs were calculated as the average of five independent regions of interest in the left ventricular area. DMD^mdx^ mice exhibited prolonged APD_90_, APD_50_, and APD_30_ at baseline (90.74 ± 5.54 ms, 39.71 ± 3.46 ms, and 19.27 ± 2.86, respectively) and in the presence of isoproterenol (77.61 ± 1.90 ms, 35.39 ± 2.24 ms, and 18.11 ± 2.54 ms, respectively) compared to wild-type mice (baseline_APD90, APD50, and APD30_: 55.05 ± 5.85 ms, 21.41 ± 2.67 ms, and 10.71 ± 0.76 ms, respectively; isoproterenol_APD90, APD50, and APD30_:51.17 ±2.83 ms, 22.64 ± 1.50 ms, and 12.32 ± 1.13 ms, respectively) (Figure 3C). Notably, DMD^mdx^:C271S^+/-^ mice demonstrated significantly shorter action potential durations, resembling those of wild-type hearts under both baseline conditions and isoproterenol stimulation (baseline_APD90, APD50, and APD30_: 55.08 ± 4.12 ms, 20.96 ± 2.19 ms, and 10.40 ± 0.35 ms, respectively; isoproterenol_APD90, APD50, and APD30_: 51.91 ± 4.27 ms, 15.06 ± 0.55 ms, and 8.75 ± 0.55 ms, respectively) (Figure 3C). These results suggest that S-nitrosylation of Cx43 hemichannels at cysteine 271 is critical for inducing prolonged action potentials in DMD^mdx^ hearts under cardiac stress.

**Figure 3.**
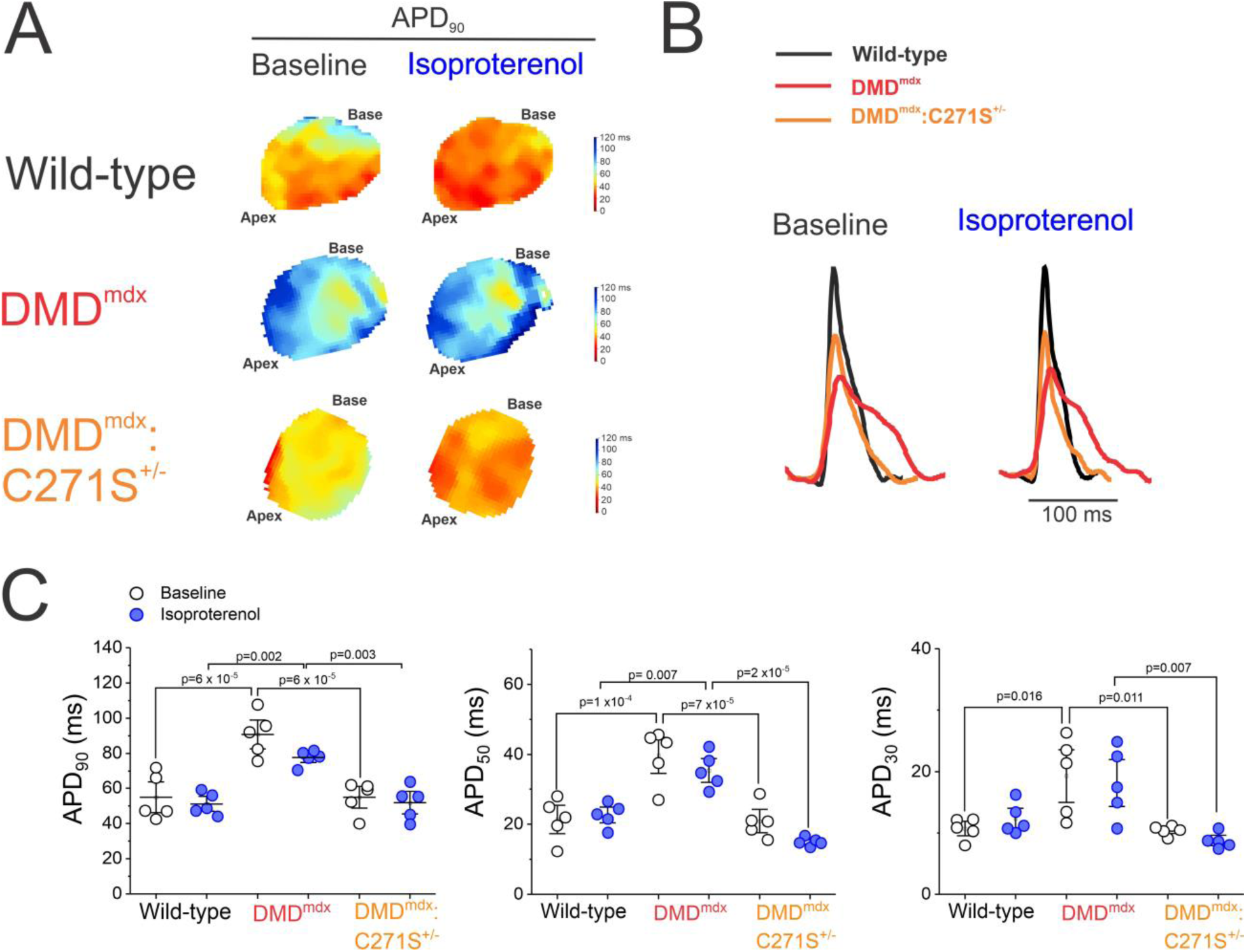
Decreased levels of S-nitrosylated Cx43 hemichannels in DMD^mdx^ mice prevent the prolongation of action potential durations (APDs) observed in DMD^mdx^ mice. (A) Representative heat maps of action potential duration at 90% repolarization (APD_90_) of wild-type, DMD^mdx^, and DMD^mdx^:C271S^+/-^ hearts before (baseline) and after isoproterenol treatment (4 µM). (B) Representative optical mapping traces of APDs at baseline conditions and following β-adrenergic stimulation (isoproterenol) in wild-type (black), DMD^mdx^ (red), and DMD^mdx^:C271S^+/-^ (orange) hearts previously loaded with the voltage-sensitive probe RH237 via Langendorff perfusion system. (C) Action potential durations: APD_90_, APD_50_, APD_30_ before (white circles) and after isoproterenol stimulation (blue circles). APDs were calculated by the average of 5 independent regions of interest in the ventricle area of the mouse heart. *n*=5 mice per group. Each dot represents an independent animal. Group comparisons were made using a one-way ANOVA followed by Tukey’s post hoc test. Data are presented as means ± SEM.

Pathological opening of Cx43 hemichannels has recently been shown to disrupt intracellular Ca^2+^ handling, promoting Ca^2+^ overload in ventricular cardiomyocytes (21, 29, 30). To investigate whether cardiac Ca^2+^ signaling depends on S-nitrosylation of Cx43, which is known to activate Cx43 hemichannels, we employed whole-heart optical mapping recordings of Ca^2+^ signals using Rhod-2AM. We evaluated Ca^2+^ signals in the presence or absence of isoproterenol (4 µM) during sinus rhythm in Langendorff-perfused hearts in the left ventricles at baseline and under isoproterenol stimulation in wild-type, DMD^mdx^, and DMD^mdx^:C271S^+/-^ mice. Figures 4A and 4B show representative heat maps of Ca^2+^ amplitudes and transient traces for wild-type, DMD^mdx^, and DMD^mdx^:C271S^+/-^ mice before and after isoproterenol stimulation. DMD^mdx^ mice exhibited significantly larger Ca^2+^ transient amplitudes and tau ratios (Ca^2+^ amplitude: 2.03 ± 0.23 a.u, tau: 1.56 ± 0.19 a.u) before and after β-adrenergic stimulation (4 µM isoproterenol) compared to wild-type (Ca^2+^ amplitude: 1.18 ± 0.08 a.u, tau: 0.66 ± 0.13 a.u) and DMD^mdx^:C271S^+/-^ hearts (Ca^2+^ amplitude: 1.20 ± 0.09 a.u, tau: 0.74 ± 0.09 a.u) (Figure 4C). These findings support the notion that S-nitrosylation of Cx43 hemichannels is crucial for intracellular cardiac Ca^2+^ handling following cardiac stress in DMD^mdx^ mice and may play a critical role in the development of cardiomyopathies.

**Figure 4.**
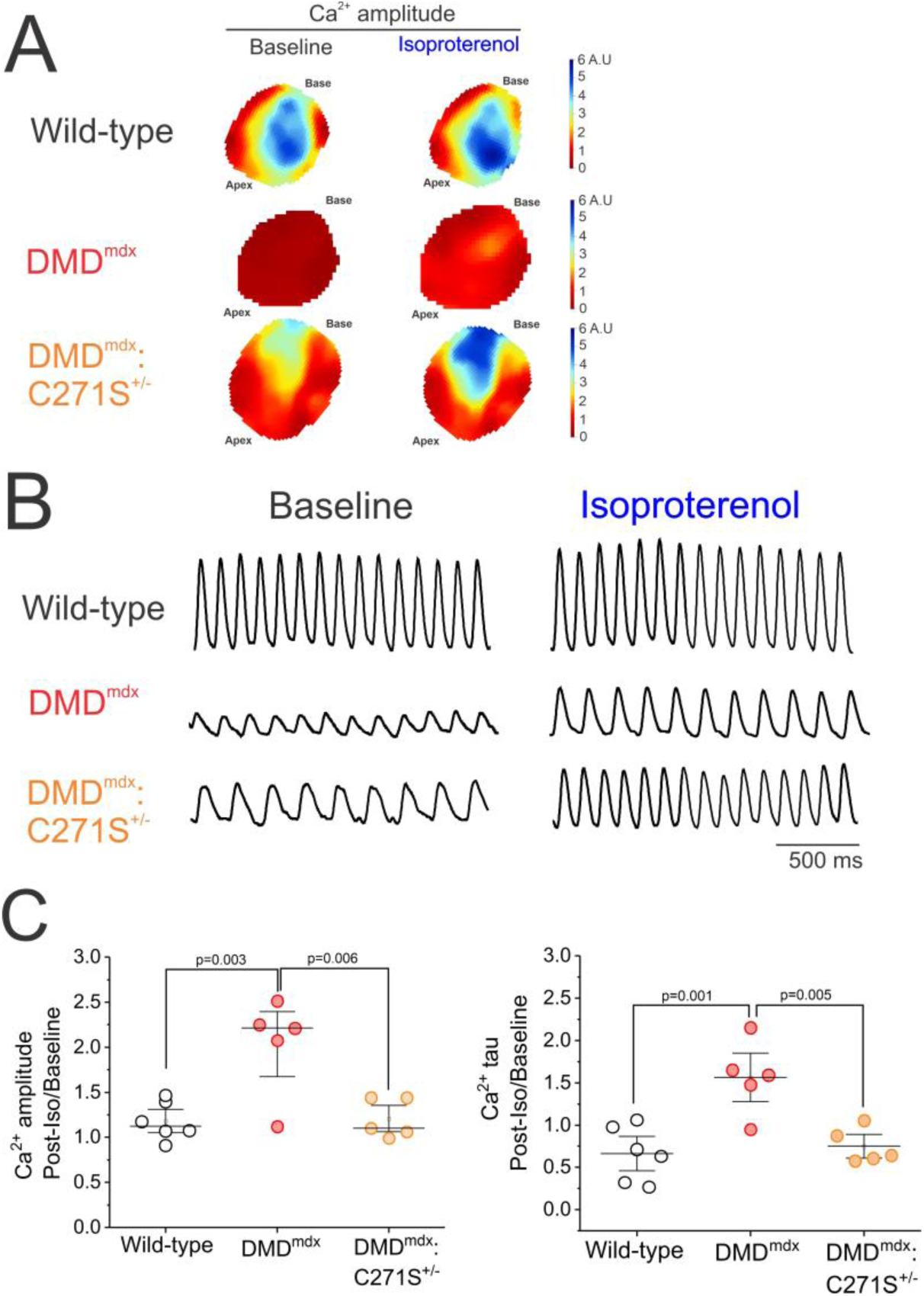
S-nitrosylation of Cx43 hemichannels in DMD^mdx^ mice mediates increased cardiac Ca^2+^ signaling. (A) Representative heat maps of Ca^2+^ amplitudes of wild-type, DMD^mdx^, and DMD^mdx^:C271S^+/-^ hearts before (baseline) and after isoproterenol treatment (4 µM). (B) Representative Ca^2+^ traces at baseline conditions and following β-adrenergic stimulation (4 µM, isoproterenol) in wild-type (black), DMD^mdx^ (red), and DMD^mdx^:C271S^+/-^ (orange) in hearts previously loaded with the Ca^2+^ indicator Rhod-2 via Langendorff perfusion system. (C) Quantification of Ca^2+^ amplitude (left panel) and decay time (right panel) ratio before and post-β-adrenergic stimulation with isoproterenol. Ca^2+^ amplitudes were calculated as the average of 5 independent regions of interest in the ventricular area of the mouse heart. *n*=6-5 mice per group. Each dot represents an independent animal. Group comparisons were made using a one-way ANOVA followed by Tukey’s post hoc test. Data are presented as means ± SEM.

### S-nitrosylation of cardiac Cx43 hemichannels in DMD^mdx^ induces action potential heterogeneity in response to frequency and β-adrenergic stimulation

We further examined the effects of repeated electrical stimulation on ventricular action potentials and Ca^2+^ signaling in whole-heart preparation of wild-type, DMD^mdx^, and DMD^mdx^:C271S^+/-^ mice. Under pathological conditions, such stimulation might lead to arrhythmogenic action potentials and Ca^2+^ transients. To investigate the role of S-nitrosylated Cx43 hemichannels, we employed optical mapping to assess both sinus rhythm and repeated electrical stimulation at the ventricular apex with a frequency of 10 Hz along with a 4 µM isoproterenol infusion, followed by a 2-minute post-stimulation recovery period. Figure 5A shows representative images of action potentials duration (APD) and Ca^2+^ transients for wild-type, DMD^mdx^, and DMD^mdx^:C271S^+/-^ hearts during sinus rhythm, 10 Hz stimulation with isoproterenol, and 2 minutes post-stimulation. Wild-type and DMD^mdx^:C271S^+/-^ hearts exhibited consistent rhythmicity in APDs and Ca^2+^ signals during sinus rhythm, electrical stimulation with β-adrenergic activation, and the 2-minute recovery phase. In contrast, DMD^mdx^ hearts displayed reduced signal amplitudes in both APDs and Ca^2+^ transients at sinus rhythm, as well as altered responsiveness to electrical and isoproterenol stimulation showing increased frequency and amplitude of signals. Post-stimulation, DMD^mdx^ hearts showed significant heterogeneity in APDs and Ca^2+^ signals. To quantify these observations, we calculated the coefficient of variation for diastolic time across beats (Figure 5B). In DMD^mdx^ hearts, the percentage of the coefficient of variation after 2 minutes of repeated stimulation with β-adrenergic activation (18.17 ± 3.83 %) was significantly higher compared to baseline (2.42 ± 1.68%) and during stimulation (3.25 ± 2.53 %). Conversely, wild-type and DMD^mdx^:C271S^+/-^ hearts displayed similar coefficients of variation post-stimulation (3.67 ± 2.08 % and 5.21 ± 2.97 %, respectively), indicating a comparable degree of heterogeneity across both conditions. Additionally, we assessed Ca^2+^ signals in the presence or absence of isoproterenol (4 µM) and electrical stimulation in wild-type, DMD^mdx^, and DMD^mdx^:C271S^+/-^ mice. Figure 5C shows that DMD^mdx^ mice exhibited significantly larger Ca^2+^ transient amplitudes (1.47 ± 0.11 a. u) after stimulation compared to wild-type (1.11 ± 0.05 a. u) and DMD^mdx^:C271S^+/-^ hearts (1.02 ± 0.06 a. u). These findings support the notion that isoproterenol and electrical stimulation-mediated activation of S-nitrosylated Cx43 hemichannels is fundamental for the heterogeneity of APDs and impaired intracellular cardiac Ca^2+^ handling in DMD^mdx^ mice.

**Figure 5.**
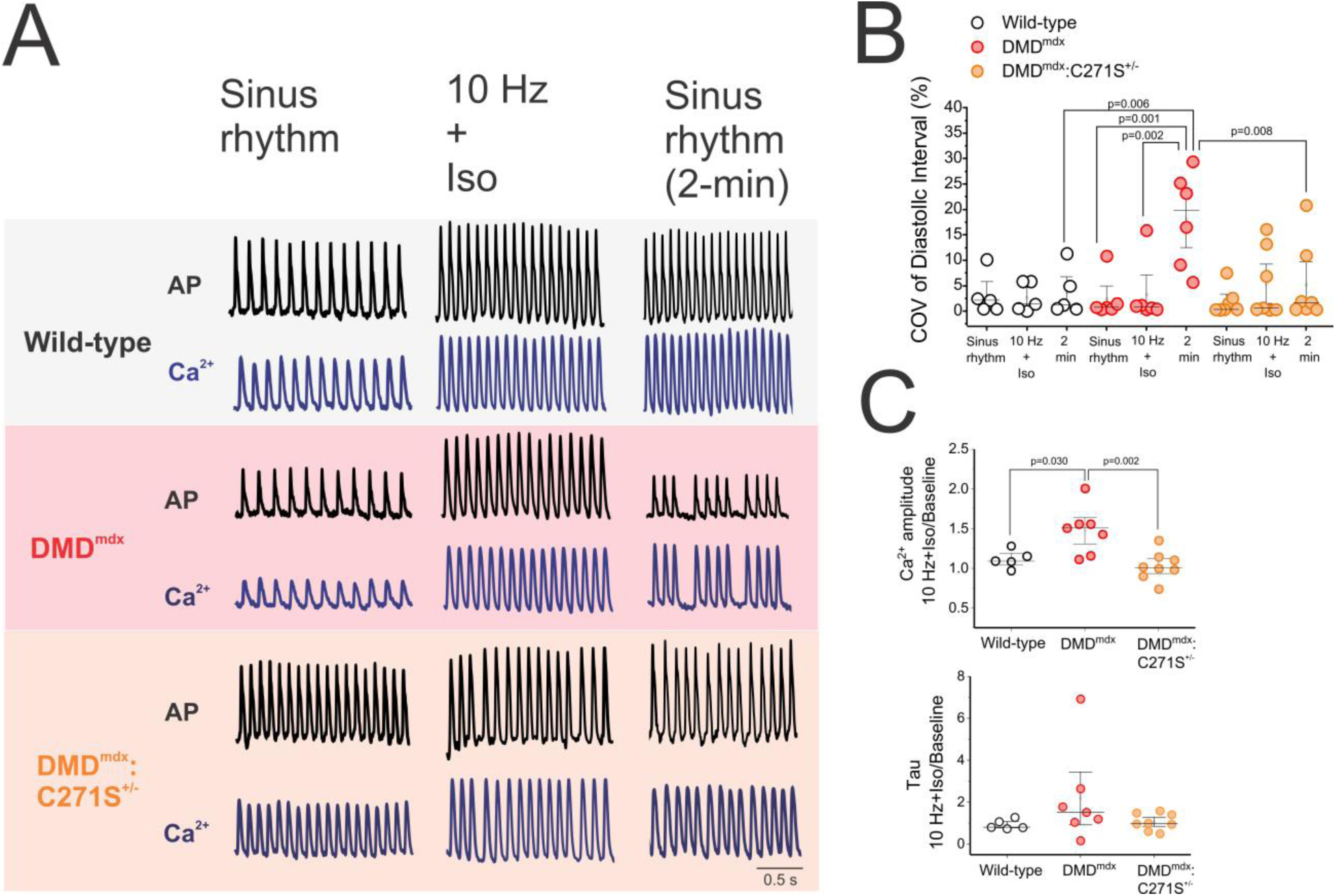
S-nitrosylation of cardiac Cx43 hemichannels in DMD^mdx^ mice mediates Ca^2+^ and action potential heterogeneity via frequency stimulation and β-adrenergic-mediated cardiac stress. (A) Representative optical mapping traces of action potentials (AP, top traces, black) and Ca^2+^ signals (bottom traces, blue) in wild-type (black), DMD^mdx^ (red), and DMD^mdx^:C271S^+/-^ (orange) hearts at sinus rhythm (left), during (middle), and 2 minutes (right) after 10 Hz burst pacing for 1 minute + 4 μM of isoproterenol (Iso) (10 Hz + Iso) stimulation. Scale bar: 0.5 s. (B) Average coefficient of variation (COV, as a percentage) of diastolic intervals between cardiac action potentials at sinus rhythm, 10 Hz + Iso, and 2 minutes after electrical stimulation in wild-type (white circles), DMD^mdx^ (red circles), and DMD^mdx^:C271S^+/-^ (orange circles) hearts. The diastolic intervals were obtained from diastolic time intervals among APs, calculated as the average of 6 consecutive beats in 6 independent regions of interest in the ventricular area of the mouse heart. (C) Quantification of Ca^2+^ amplitude (top panel) and decay time (right panel) ratio before and post-β-adrenergic stimulation with isoproterenol (Iso) + 10 Hz. Ca^2+^ amplitudes and tau were calculated as the average of 5 independent regions of interest in the ventricular area of the mouse heart. n=5-7 mice per group. Each dot represents an independent animal. Group comparisons were made using a one-way ANOVA followed by Tukey’s post hoc test. Data are presented as means ± SEM.

### DMD^mdx^:C271S^+/-^ mice are protected against myocardial injury induced by isoproterenol stimulation

Several cases of myocardial injury have been reported in patients with DMD, but the underlying etiological mechanisms remain poorly understood (4–7). Both DMD patients and DMD mouse models exhibit elevated levels of cardiac troponin I (cTnI), which is significantly associated with myocardial injury and the development of cardiomyopathies throughout life (6, 34–37). In line with this idea, highly sensitive cardiac troponin assays are employed to assess acute myocardial injury in humans (38, 39). To investigate the hypothesis that S-nitrosylation of Cx43 is involved in myocardial injury, we examined whether decreased levels of S-nitrosylated hemichannels prevent elevated cTnI levels in DMD^mdx^ mice under cardiac stress, using a high-sensitivity cTnI ELISA Kit (Figure 6). Plasma levels of cTnI significantly increased 4 hours post-isoproterenol treatment in DMD^mdx^ mice (7.83 ± 2.05 ng/ml) compared to baseline conditions (0.34 ± 0.06 ng/ml). Upon β-adrenergic stimulation, DMD^mdx^ mice treated with Gap19 (1.15 ± 0.22 ng/ml), DMD^mdx^:C271S^+/-^ (2.20 ± 0.82 ng/ml), C271S^+/-^ (0.79 ± 0.42 ng/ml), and wild-type mice (0.40 ± 0.08 ng/ml) showed considerably lower levels of plasma cTnI, similar to the baseline levels of each experimental condition (Figure 6A). Following 24-h of isoproterenol stimulation, myocardial injury was assessed using the 2,3,5-Triphenyltetrazolium chloride (TTC) staining in 2-mm ventricular heart sections to evaluate whether S-nitrosylated Cx43 hemichannels are involved in ventricular myocardial injury in DMD^mdx^ mice under β-adrenergic stimulation. Isoproterenol-treated hearts from DMD^mdx^ (41.33 ± 5.85%) and Src-Gap19 (41.54 ± 1.91%) mice exhibited larger areas of myocardial injury compared to wild-type conditions (7.17±1.58%) after 24-h post-cardiac stress when compared. Strikingly, there were no significant differences in infarct areas between DMD^mdx^ mice previously treated retro-orbitally with the Cx43 hemichannel blocker, Gap19 (10 mg/kg) (15.38 ± 2.62%), DMD^mdx^:C271S^+/-^ (15.90 ± 3.36%), C271S^+/-^ (13.40 ± 5.87%), and wild-type mice (7.17 ± 1.58%) (Figure 6B and Figure 6C). Consistent with these results, DMD^mdx^ mice exhibited ∼40% mortality after 24 hours post-isoproterenol stimulation, which was prevented in DMD^mdx^:C271S^+/-^ mice, indicating that reduced levels of S-nitrosylation of Cx43 are sufficient to rescue the survival percentage observed in wild-type conditions (100% survival in mouse models of DMD^mdx^:C271S^+/-^ knock-in colonies) (Figure 6D). These findings indicate that lateralized S-nitrosylation of Cx43 hemichannels critically contributes to myocardial injury following β-adrenergic receptor activation in DMD^mdx^ mice.

**Figure 6.**
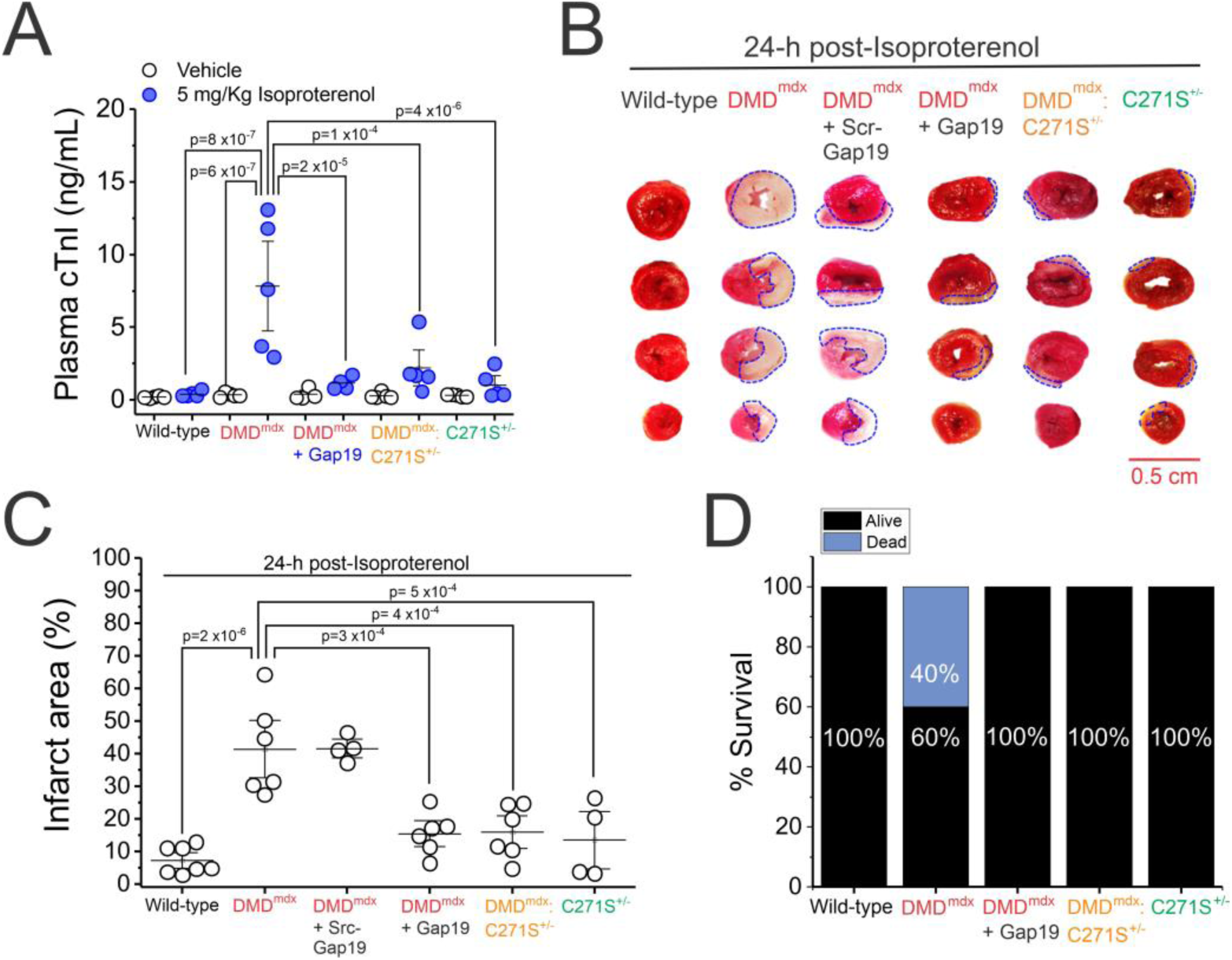
Reduced S-nitrosylated cardiac Cx43 hemichannels attenuate myocardial injury after β-adrenergic stress. (A) Plasma cardiac troponin I (cTnI) levels 4 hours after isoproterenol-mediated cardiac stress in wild-type, DMD^mdx^, DMD^mdx^ treated with Gap19 via retroorbital injection (10 mg/kg), DMD^mdx^:C271S^+/-^, and C271S^+/-^ mice. (B) Representative images of cross-sections of mouse ventricles stained with TTC 24 hours post-isoproterenol stimulation. Myocardial injury was outlined by a blue line. Scale bar: 0.5 cm. (C) Quantification of % + TTC area of heart ventricles after β-adrenergic stimulation with respect to the total slice areas in wild-type, DMD^mdx^, DMD^mdx^ previously treated with Gap19 via retroorbital injection (10 mg/kg), DMD^mdx^ treated with Scramble-Gap19 (Src-Gap19, sequence: IKFKKEIKQ, 10 mg/kg) via retroorbital injection (10 mg/kg), DMD^mdx^:C271S^+/-^, and C271S^+/-^ mice. (D) Percentage of survival of wild-type, DMD^mdx^, DMD^mdx^ treated with Gap19 via retroorbital injection (10 mg/kg), DMD^mdx^:C271S^+/-^, and C271S^+/-^ mice 24 hours post-isoproterenol stimulation. Each dot represents an independent mouse. *n*=4-7 mice per group. Group comparisons were made using a one-way ANOVA followed by Tukey’s post hoc test. Data are presented as means ± SEM.

## DISCUSSION

While we have consistently demonstrated a crucial pathological role for lateralized Cx43 hemichannels in DMD cardiomyopathies (18–20), the mechanisms associated with the hyperactivity of Cx43 hemichannels remain unsolved (19). In this study, we evaluated the pathological role of S-nitrosylation in Cx43 at cysteine 271 evoking deadly arrhythmias and myocardial injury under β-adrenergic-evoked cardiac stress. We found that β-adrenergic stimulation in DMD^mdx^ provokes severe arrhythmias, aberrant Ca^2+^ signaling, prolonged action potential duration, and myocardial injury but these pathological features are prevented in DMD^mdx^:C271S^+/-^ mice with impaired Cx43 S-nitrosylation. These findings demonstrate that S-nitrosylation of Cx43 hemichannels at cysteine 271 is critical for β-adrenergic-induced lethal arrhythmias and myocardial injury in DMD cardiomyopathy.

Duchenne muscular dystrophy (DMD) patients exhibit an imbalance in the autonomic nervous system, characterized by decreased parasympathetic activity and increased sympathetic activity (40, 41). The pathological cardiac features associated with DMD include increased heart rates, tachycardia, and sudden cardiac death (7, 40, 42). The DMD^mdx^ mouse model also displays tachycardia, decreased heart rate variability, and an imbalance in autonomic nervous system modulation, with decreased parasympathetic and increased sympathetic activity. Thus, it is a valuable experimental model for understanding DMD cardiomyopathy (42). In DMD^mdx^ mice, β-adrenergic–induced arrhythmias and sudden death are primarily mediated by the hyperactivity of lateralized Cx43 hemichannels (18, 19). While the mechanisms underlying lateralization have been partially unveiled, the opening of hyperactive Cx43 hemichannels remains less understood. In particular, Cx43 remodeling has been long associated with cardiac dysfunction in models of heart failure (43–45), myocarditis (46), ischemia (17, 47), and DMD^mdx^ cardiomyopathy (19, 20). This lateralization results from reduced post-translational phosphorylation at C-terminal serine S325/S328/S330 in human and mouse DMD^mdx^ hearts via a microtubule-dependent reorganization mechanism (18, 48). Importantly, DMD^mdx^:C271S^+/-^ mice, which have impaired Cx43 S-nitrosylation, displayed similar Cx43 protein lateralization as observed in DMD^mdx^ hearts sections. This result indicates that S-nitrosylation is not a post-translational modification involved in Cx43 cardiac lateralization. Therefore, the cardiac protection observed in DMD^mdx^:C271S^+/-^ mice is likely due to the reduced or blunted opening of Cx43 hemichannels rather than the prevention of remodeling.

We previously showed that treating DMD^mdx^ mice with a non-selective NOS inhibitor, L-NAME, results in reduced levels of S-nitrosylation of Cx43 (19). This treatment also led to decreased membrane permeability and mitigated excitability disturbances preventing Cx43-mediated delayed afterdepolarizations (DADs) and triggered activity (TA) in isolated cardiomyocytes, thereby lowering the severity of arrhythmogenicity (19). These results are in line with our current data showing that isoproterenol-induced arrhythmias are significantly lower in DMD^mdx^:C271S^+/-^ when compared to DMD^mdx^ mice, supporting a specific role for S-nitrosylated Cx43 hemichannels in the development of life-threatening arrhythmias. This study represents the first to suggest that C271 in Cx43 is a key amino acid residue in regulating Cx43-mediated cardiac hemichannel currents, which could be an important therapeutic target for future studies in cardiac pathology.

In our whole-heart optical mapping experiments, we consistently found that DMD^mdx^ mice exhibited prolonged action potentials at baseline, which remained extended in the presence of isoproterenol, whereas no such prolongation was observed in DMD^mdx^:C271S^+/-^ hearts (Figure 3). Strikingly, DMD^mdx^:C271S^+/-^ hearts displayed shorter action potential durations under baseline conditions and in response to isoproterenol, similar to wild-type conditions (Figure 3). The action potential plateau and early repolarization in a mouse action potential, represented by APD_30_ and APD_50_, are characterized by rapid outward K^+^ currents that account for more than 50% of the total outward K^+^ currents in left ventricular epicardial myocytes (49, 50). This indicates that S-nitrosylation of Cx43 plays a pivotal role in regulating cardiac excitability during both early (APD_30_ and APD_50_) and late phases of repolarization, including APD_90_. S-nitrosylation of Cx43 hemichannels in a whole-heart preparation appears to influence local membrane potentials, helping to maintain prolonged action potentials during β-adrenergic stimulation without extending them beyond baseline levels. These electrophysiological parameters could serve as early markers of cardiac pathology during various stages of the repolarization phase of the cardiac action potential.

Dystrophic cardiomyocytes also exhibit abnormal intracellular Ca^2+^ signaling (51); however, the mechanisms underlying this dysregulation are not yet fully understood (29). Consistently, our data showed that DMD^mdx^ hearts show increased Ca^2+^ levels in the left ventricle during optical mapping experiments following β-adrenergic stimulation with isoproterenol. One proposed mechanism for Ca^2+^ mishandling involves structural and functional remodeling of ryanodine receptors in the DMD^mdx^ model. Leaky RyR2 and diastolic sarcoplasmic reticulum Ca^2+^ leak have been shown to mediate aberrant depolarization of isolated cardiomyocytes, leading to arrhythmias *in vivo* (51). Interestingly, Cx43 colocalizes with large dyadic RyR superclusters, forming microdomains at the perinexus. It has been also reported that Ca^2+^ release from the sarcoplasmic reticulum opens Cx43 hemichannels in isolated mouse and pig left ventricular cardiomyocytes under wild-type conditions, as well as in human heart failure (29). Consistently, DMD^mdx^:C271S^+/-^ hearts show only a slight increase in Ca^2+^ amplitude and similar Tau ratios to those found in wild-type hearts. This supports the idea that S-nitrosylation of Cx43 hemichannels mediates Ca^2+^ dysfunction in dystrophic cardiomyocytes.

Interestingly, the activation of Cx43 hemichannels has been suggested to be modulated by frequency and β-adrenergic stimulation, similar to other channels such as RyRs, which are dependent on Ca^2+^/calmodulin-dependent kinase II (CaMKII) activation during high-frequency stimulation at the dyad in the T-tubules (29, 52, 53). The open probability of Cx43 channels has been shown to increase at frequencies of 0.5 Hz and 4 Hz in mouse and pig ventricular cardiomyocytes (29). Correspondingly, our data indicate that increased frequency stimulation approaching 10 Hz, combined with isoproterenol stimulation, enhances action potential heterogeneity and the amplitude of intracellular Ca^2+^ signaling, thus augmenting the heterogeneity of diastolic time among action potentials on a beat-to-beat basis (Figure 5). Our results reveal that the coefficient of variation in diastolic time does not increase during high-frequency stimulation but does so during the recovery period, 2 minutes post high-frequency and adrenergic stimulation (Figure 5). This suggests that the recovery phase after tachyarrhythmic events plays a crucial role in mediating the activation of S-nitrosylated Cx43 hemichannels, thereby contributing to the disruptions in Ca^2+^ homeostasis and action potentials, and to the generation of life-threatening arrhythmias.

Multiple reports have documented distinctive signs of myocardial injury in DMD patients (4, 5, 54), such as pathological QRS complexes (55, 56), elevated cardiac troponin I (cTnI) levels in the blood (6, 34, 57), and elevated ST segments (56, 58). However, the etiological mechanisms underlying these clinical manifestations remain unclear. Importantly, Cx43 remodeling has been implicated in myocardial infarction and arrhythmia susceptibility (59). Recent studies have identified four missense variants in GJA1, the gene encoding Cx43, in patients experiencing severe arrhythmias (19), which support the involvement of Cx43 in arrhythmogenicity. In this study, for the first time, we explored the role of cardiac Cx43 hemichannels in myocardial injury within the context of DMD cardiomyopathies. Our data reveal that 24 hours post-isoproterenol challenge, DMD^mdx^ mice exhibit elevated plasma concentrations of cTnI, mirroring the increases reported in DMD humans and mouse models following myocardial injury (11). Remarkably, this elevation was abolished in DMD^mdx^ previously treated with the Cx43 hemichannel blocker Gap19 and in DMD^mdx^:C271S^+/-^ mice, strongly suggesting that S-nitrosylation and activation of Cx43 hemichannels mediate myocardial injury. Additionally, our findings indicate that 24 hours post-isoproterenol stimulation, DMD^mdx^ mice display significantly larger infarct areas. However, the infarct size was effectively reduced in DMD^mdx^ mice treated with Gap19 and in DMD^mdx^:C271S^+/-^ mice. These results highlight the pivotal role of S-nitrosylation of Cx43 hemichannels in exacerbating myocardial injury in DMD cardiomyopathy and highlight potential therapeutic avenues targeting Cx43 to mitigate cardiac damage in this disease.

In conclusion, our findings provide compelling evidence that isoproterenol-induced S-nitrosylation of Cx43 hemichannels at cysteine 271 is crucial for the hyperactivity of lateralized Cx43 hemichannels in the cardiac ventricle. This post-translational modification significantly augments Ca^2+^ signaling and prolongs action potentials, leading to fatal arrhythmias and myocardial injury in DMD mice. These results underscore the significance of understanding the underlying mechanisms of Cx43 hemichannel modulation, positioning post-translational modifications of Cx43 as promising therapeutic targets for addressing lethal arrhythmias and myocardial injury.

## Material and Methods

### Animals

DMD^mdx^ mice (4-6 months old) were obtained from Jackson Laboratory. Mice were heparinized (5,000 U/kg) and then overdosed with isoflurane. We recently published that the S-nitrosylation of Cx43 occurs specifically at cysteine 271 (C271) at the C-terminal of Cx43 (19). Consistent with this finding, we generated a knock-in mouse line using CRISPR-Cas9 genome editing technology, in which the site of S-nitrosylation of Cx43 is substituted by serine (C271S). After generating stable colonies of heterozygous C271S^+/-^ and homozygous C271S^+/+^ mice, normal growth, breeding, and survival were observed. Finally, DMD^mdx^ mice were crossed to C271S colonies, obtaining DMD^mdx^:C271S^+/-^ mice. Genotyping was performed using Transnetyx. Mice were provided with food and water ad libitum. All experiments involving mice were approved by the Institutional Animal Care and Use Committee (IACUC) of the School of Medicine at the University of California, Davis, and performed in accordance with the National Institutes of Health guidelines.

### Immunofluorescence in cryosections of mouse heart ventricles

Mouse ventricle hearts were quick-frozen in liquid nitrogen and stored at −80°C. 6 µm heart ventricle sections were obtained from cryosectioning. Samples were thawed at room temperature (RT) and washed three times with 1x PBS. Heart samples were permeabilized with 0.5% Triton X-100 in PBS for 30 minutes and washed 3 times with PBSt (1x PBS + 0.1% Tween 20). Subsequently, samples were incubated with blocking buffer, 10% normal donkey serum, for 1 hour at RT (#005-000-121, Jackson ImmunoResearch). Sections were incubated with Cx43 (1:1,000 dilution, anti-rabbit, #C6219, Sigma-Aldrich) and N-cadherin (1:100 dilution, anti-mouse, #33-3900, Invitrogen) antibodies in blocking buffer at 4°C overnight. Heart sections were washed with PBSt and then incubated with goat Alexa Fluor secondary antibodies (Jackson ImmunoResearch). Sections were washed with PBSt, and incubated with Wheat Germ Agglutinin (WGA), Alexa Fluor 350 (#W11263), for 1 hour to mark the cell membrane of cardiomyocytes. Lastly, coverslips were mounted on heart sections using ProLong^TM^ Gold antifade reagent (#P36930, Invitrogen). Images were taken on a Zeiss LSM880 Airyscan confocal microscope and analyzed using the software ImageJ.

### Electrocardiogram telemetry

Mice were anesthetized with 4% isoflurane for induction, followed by 2% for maintenance. Electrocardiograms (ECGs) were performed using lead II via implantation of a wireless radio telemeter subcutaneously (ETA-F10 from DSI). Buprenorphine (0.05-0.1 mg/kg) was injected subcutaneously after the surgery and then as needed if signs of pain were present. Mice recovered for 3 days, and on the 4th day post-electrode implantation, they were first weighed and then separated into single cages placed on telemetry receivers. Mice were treated with isoproterenol (5 mg/kg i.p.; no more than 0.25 mL; #1351005, USP), and ECGs were continuously recorded and evaluated for 24 hours post-injection. A 1-hour baseline reading was taken to monitor activity and obtain ECG recordings. All cardiac parameters were measured using the ER400 energizer/receiver, and data were collected using Ponemah software. The total amount of arrhythmias included single premature ventricular contractions (PVCs), double PVCs, and three or more PVCs (non-sustained ventricular tachycardia, NSVT).

### Ca^2+^ transients and ΔV_m_ changes in optical mapping in the intact mouse heart

The mice were anesthetized using isoflurane and euthanized by cervical dislocation. The hearts were rapidly isolated and then Langendorff-perfused through the aorta with a warmed and oxygenated KH solution (in mM: 119 NaCl, 25 NaHCO₃, 4 KCl, 1.2 KH₂PO₄, 1 MgCl₂, 1.8 CaCl₂·2H₂O, 10 D-glucose) at a flow rate of 3 mL/minute, maintaining a temperature of 37°C with a gas mixture of 95% O₂ and 5% CO₂. Once the heart was cannulated, it was placed in the optical mapping setup (MappingLab, UK). Optical mapping recordings were captured using two high-speed sCMOS cameras (Prime BSI Express, Teledyne Photometrics) with high quantum efficiency. For imaging, heart preparations were excited at 530 ± 20 nm using a light-emitting diode (LED) epi-illumination. The emission light passing through a premium long-pass filter (FELH550) was split using a dichroic mirror (Thorlabs; DMLP638). The emission light was then passed through a 700 nm long-pass filter (FELH700) and a 585 nm band-pass filter (Thorlabs; FBH585) for ΔVm and Ca²⁺ imaging, respectively. To prevent motion artifacts, hearts were mechanically uncoupled using 4 µM blebbistatin (#TRC-B592500, TRC) and 0.008% Pluronic acid F-127 (#540025, Millipore) following a previously described protocol (60). Simultaneous recording of ΔVm and Ca²⁺ transients was performed after loading the hearts with RH237 (N-(4-Sulfobutyl)-4-(6-(4-(Dibutylamino)phenyl)hexatrienyl)Pyridinium, Inner Salt, #S1109, ThermoFisher) and Rhod-2AM (#R1244, ThermoFisher), respectively. Pharmacological agonists were introduced directly into the perfusion after obtaining baseline recordings, and the recordings were evaluated 4 minutes after the injection of isoproterenol during normal sinus rhythm. For experiments with frequency stimulation, ΔVm and Ca²⁺ transients were recorded under baseline conditions, during stimulation together with 4 µM isoproterenol, and 2 minutes post-stimulation at sinus rhythm, representing the recovery time. During stimulation, electrodes were strategically positioned in the ventricular region of the heart, and hearts were stimulated at 10 Hz at a constant current of 1 nA for 1 minute. Ca²⁺ amplitudes and APDs were calculated as the average of 5 independent regions of interest in the ventricular area of the mouse heart. The coefficient of variation between APDs was measured as the standard deviation/average of the diastolic intervals. The diastolic intervals were obtained from diastolic time intervals among APs, calculated as the average of 6 consecutive beats in 6 independent regions of interest in the ventricular area of the mouse heart. Fluorescence images were captured at a sample rate of 1200 fps, with a region size of 150 x 150 pixels and an exposure time of 0.7 seconds. Optical mapping data was acquired in real-time using an 8-channel TTL analog-to-digital converter with OMapRecord 5.0 software and processed with OMapScope5.9.4 (Mapping Lab).

### Measurement of plasma cardiac troponin I (cTnI)

Blood was collected from the tail vein of isoflurane-anesthetized mice before and 4 hours after the isoproterenol injection. The blood samples were stored in EDTA-containing tubes and kept on ice. Following centrifugation at 4,000 RPM for 20 minutes, plasma samples were stored at -80°C until ELISA analysis. Serum cardiac troponin I levels were measured using the high-sensitivity mouse cardiac Troponin-I ELISA Kit (#CTNI-1-HSP, Life Diagnostics, Inc.) on the multimode plate reader, VICTOR Nivo (PerkinElmer Life).

### 2,3,5-triphenyltetrazolium chloride (TTC) staining

TTC staining was performed to evaluate myocardial injury (61). After heart isolation, hearts were kept at -20°C for 1 h. Using a razor blade, the hearts were cut transversely into 2-mm-thick slices. The slices were then incubated in 1% TTC in 1x PBS for 1 hour at 37°C, followed by incubation in 4% paraformaldehyde (PFA) for 30 minutes at room temperature (RT). Finally, the slices were transferred to 1x PBS. Pictures were taken using a stereoscopic microscope coupled with a camera at 20x magnification.

### Molecular biology

cDNA for human Cx43 was synthesized and subcloned into a pGEM-HA vector. The single mutation C271S in Cx43 was introduced using the QuickChange II Site-Directed Mutagenesis Kit (Agilent Technologies) and confirmed by Sanger sequencing. cDNAs were transcribed in vitro to cRNAs using the HiScribe® T7 Arca mRNA Kit (New England BioLabs). The Cx43 antisense oligonucleotide (Sequence: 5’ GCT TTA GTA ATT CCC ATC CTG CCA TGT TTC 3’ (63) was synthesized by Integrated DNA Technologies (Coralville, IA).

### Expression of Cx43 in Xenopus oocytes

Oocytes from female *Xenopus laevis* (Xenopus 1 Corp, Dexter, MI) were collected and digested as we described previously (63, 64) in strict adherence to the protocol approved by the Institutional Animal Care and Use Committee (IACUC) at the University of California Davis and conforming to the National Institutes of Health Guide for the Care and Use of Laboratory Animals. Defolliculated oocytes (stage IV-V) were individually microinjected with 25 ng of cRNAs for human Cx43 wild-type (WT), human Cx43 C271S, or a combination of Cx43 WT plus Cx43 C271S (ratio 1:1). Oocytes were also injected with an antisense oligonucleotide (AS) to prevent the expression of endogenous Xenopus Cx38, as previously described by Ebihara (63). Non-injected oocytes or oocytes injected with AS alone were used for control experiments. After microinjection, oocytes were stored for 2 days at 16 °C in sterile plates containing ND96 solution (composition in mM: 96 NaCl, 2 KCl, 1 MgCl_2_, 1.8 CaCl_2_, 5 HEPES, adjusted to pH 7.4) supplemented with streptomycin (50 µg/mL) plus penicillin (50 Units/mL).

### Two-Electrode Voltage-Clamp (TEVC)

TEVC technique was performed as previously described by us (64). Briefly, two pulled borosilicate glass micropipettes were filled with 3 M KCl, resulting in resistances ranging from 0.2 to 2 MΩ. The electrical signal was amplified using an Oocyte Clamp Amplifier (OC-725C, Warner Instrument Corp., USA) and digitized by a data acquisition system (Digidata 1440A, Molecular Devices, USA). Data were sampled at 2 kHz, and analyzed using pClamp 10 software (Molecular Devices, USA). Experiments were conducted at room temperature (20-22 °C). The recording solution contained (in mM) 117 Tetraethylammonium chloride (TEACl), 0.2 CaCl_2_, and 5 HEPES, with the pH adjusted to 7.40. Current-voltage relationship was obtained by analyzing the magnitude of activation currents evoked by depolarizing pulses (from -80 mV to +80 mV; holding potential: -80 mV; duration of depolarization pulse: 2 s). After recording ionic currents under control conditions, oocytes were incubated with 100 µM DEA NONOate for 15 min, and ionic currents were recorded again. To confirm that currents were mediated by Cx43, we incubated the oocytes with 200 µM LaCl_3_ in the presence of DEA NONOate at the end of each experiment. Only those experiments with La^3+^-sensitive currents were considered for the analysis of the Cx43 WT group.

### Biotin-switch for S-nitrosylated proteins

To evaluate S-nitrosylation levels of Cx43 proteins in knock-in mice, we isolated S-nitrosylated Cx43 proteins from ventricular hearts obtained from wild-type, DMD^mdx^, DMD^mdx^:C271S^+/-^, C271S^+/-^, and C271S^+/+^ mice. Firstly, heart samples were homogenized in HEN buffer (250 mM HEPES, 0.1 EDTA, and 0.1 mM Neocuproine, protease inhibitor cocktail, pH=7.7). Biotin switch protocol was performed to 200 µg of protein samples from wild-type, DMD^mdx^, DMD^mdx^:C271S^+/-^ , C271S^+/-^, and C271S^+/+^ mice based on (65). Briefly, heart samples were treated with S-methyl methanethiosulfonate (#208795, Sigma Aldrich) at 50°C for 1 hour to block cysteine-free thiols. Subsequently, proteins were precipitated by incubating them in acetone at -20°C for 24 hours. Samples were washed and solubilized. Heart samples underwent incubation with Biotin-HPDP (#16459, Cayman Chemicals) and sodium L-ascorbate (#11140, Sigma Aldrich) to convert S-nitrosylated cysteine residues into free cysteine. Proteins were precipitated again to remove any excess of Biotin-HPDP and solubilized to run Western Blot. After solubilization, the heart samples were incubated with Streptavidin Agarose Resin (#20349, ThermoScientific) and then centrifuged at 24,000 RPM for 20 minutes at 4°C to pull down the respective biotin-tagged proteins. The proteins were subsequently separated via electrophoresis using 12% SDS-PAGE gels and transferred onto a PVDF membrane (Trans-Blot Turbo, Bio-Rad). S-nitrosylated Cx43 proteins were detected using a mouse anti-connexin-43 antibody (#C8093, Sigma-Aldrich, dilution: 1:8,000). ImageJ software was used to analyze the intensity of the signal.

### Measurement of arterial blood pressure

To measure arterial blood pressure, mice were anesthetized using isoflurane at a concentration of 1.5%, ensuring deep anesthesia was achieved, as indicated by the absence of reflexes. The anesthetized mice were placed on a heating pad in a supine position, and the surgical field was prepared with sterile drapes. A midline incision was performed in the neck to expose the carotid artery. A sterile catheter connected to a pressure transducer (ADInstruments, Australia) was inserted into the artery. The catheter was secured and connected to the pressure transducer, and the system was flushed with sterile or heparinized saline to remove any air bubbles. Arterial blood pressure was recorded using LabTutor (version 2012), the pressure transducer was calibrated, and proper zeroing was ensured. Throughout the data collection process, both the mouse and the catheter were continuously monitored to ensure stable measurements. After data collection was completed, the catheter was carefully removed, the incision was closed with sutures or surgical glue, and sterile eye ointment was applied. Pressure changes were assessed 30 minutes post-surgery by calculating an average of four independent measurements taken every 10 minutes.

### Chemicals

Isoproterenol Hydrochloride was obtained from USP (#1351005). (S)-(-)-Blebbistatin was purchased from LGC standard (#TRC-B592500). Lanthanum chloride and diethylamine NONOate sodium salt hydrate (DEA NONOate) were obtained from Sigma-Aldrich (#D184). Heparin was obtained from Fresenius Kabi (NDC #63323-047-01). The connexin 43 hemichannel, Gap19 (#5353), was obtained from Tocris, and the Gap19-scramble (Sequence: IKFKKEIKQ) was synthesized in LifeTein.

### Statistical analysis

The data are presented as mean ± standard error of the mean. Group comparisons were conducted using either one-way ANOVA with Tukey’s post hoc test. When two groups were analyzed, a paired two-tailed Student’s t-test was performed. A significance level of p < 0.05 was used to determine statistical significance. The p-values from statistical analyses among groups are indicated in each figure. Statistical analyses were performed using Origin Pro 9.85.

## Authors contributions

Author contributions: Manuel F. Muñoz, J. Quan, T. Nguyen, Janet Nuno, Adrian Sheehy, Pablo S. Gaete, Pia C. Burboa, Mauricio A. Lillo, and JE Contreras, designed experiments. Manuel F. Muñoz performed most of the experiments. Manuel F. Muñoz, J. Quan, T. Nguyen, Janet Nuno, Adrian Sheehy, Pablo S. Gaete, Pia C. Burboa, Mauricio A. Lillo, and J.E. Contreras analyzed the data. Manuel F. Muñoz and JE Contreras wrote the original draft manuscript. Manuel F. Muñoz and JE Contreras edited the final manuscript. All authors reviewed and approved the final draft.

## Acknowledgments

This research work was supported by the AHA post-doctoral fellowship 23POST1027462 to M.F.M and NIH grant 1R01GM099490 and R21HL163930 to J.E.C. This work was supported by an American Heart Association AHA Career Development Award 932684 to M.A. Lillo, AHA Research Supplement to Promote Diversity in Science 23DIVSUP1054931 to Pia Burboa.

