## Supplementary Figures for "Impaired S-nitrosylation of Cx43 prevents arrhythmogenicity and myocardial injury upon cardiac stress in Duchenne Muscular Dystrophy"

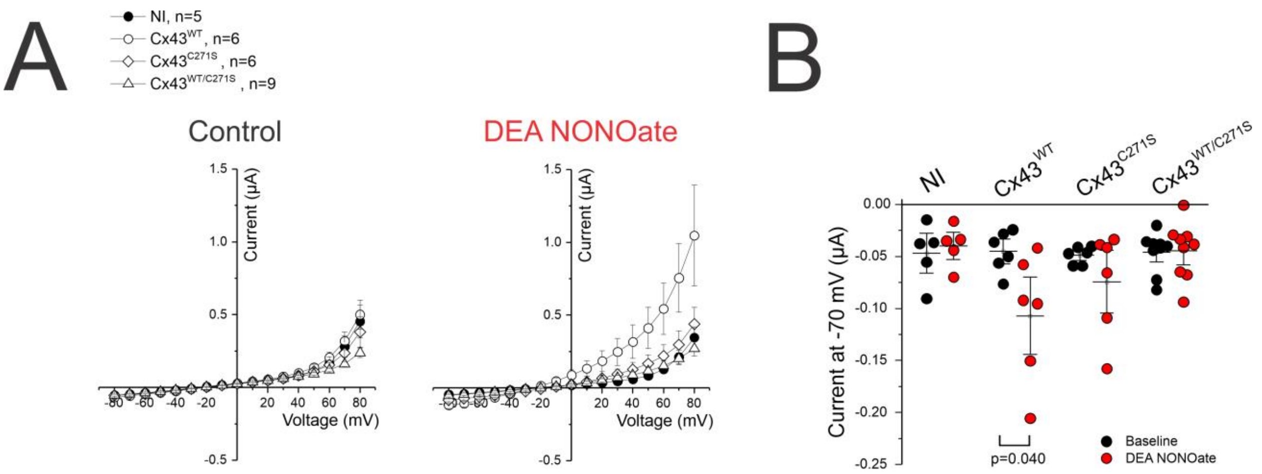


**Supplementary Figure 1. Nitric oxide (NO) donors induce hemichannel currents in oocytes expressing Cx43 hemichannels.** (A) Quantification of the current-voltage relationship in non-injected oocytes (NI) (black circles), oocytes expressing wild-type Cx43 ($Cx43^{WT}$) (white circles), the C271S mutant ($Cx43^{C271S}$) (white rhombus), and wild-type Cx43 ($Cx43^{WT}$): C271S mutant ($Cx43^{WT/C271S}$) (white triangle), under control conditions (left panel) and in the presence of 100 µM DEA NONOate (right panel). (B) Quantification of currents at -70 mV in oocytes under control conditions and in the presence of 100 µM DEA NONOate. Each dot represents an independent oocyte. Group comparisons were made using a paired Student's t-test. Data are presented as means ± SEM.

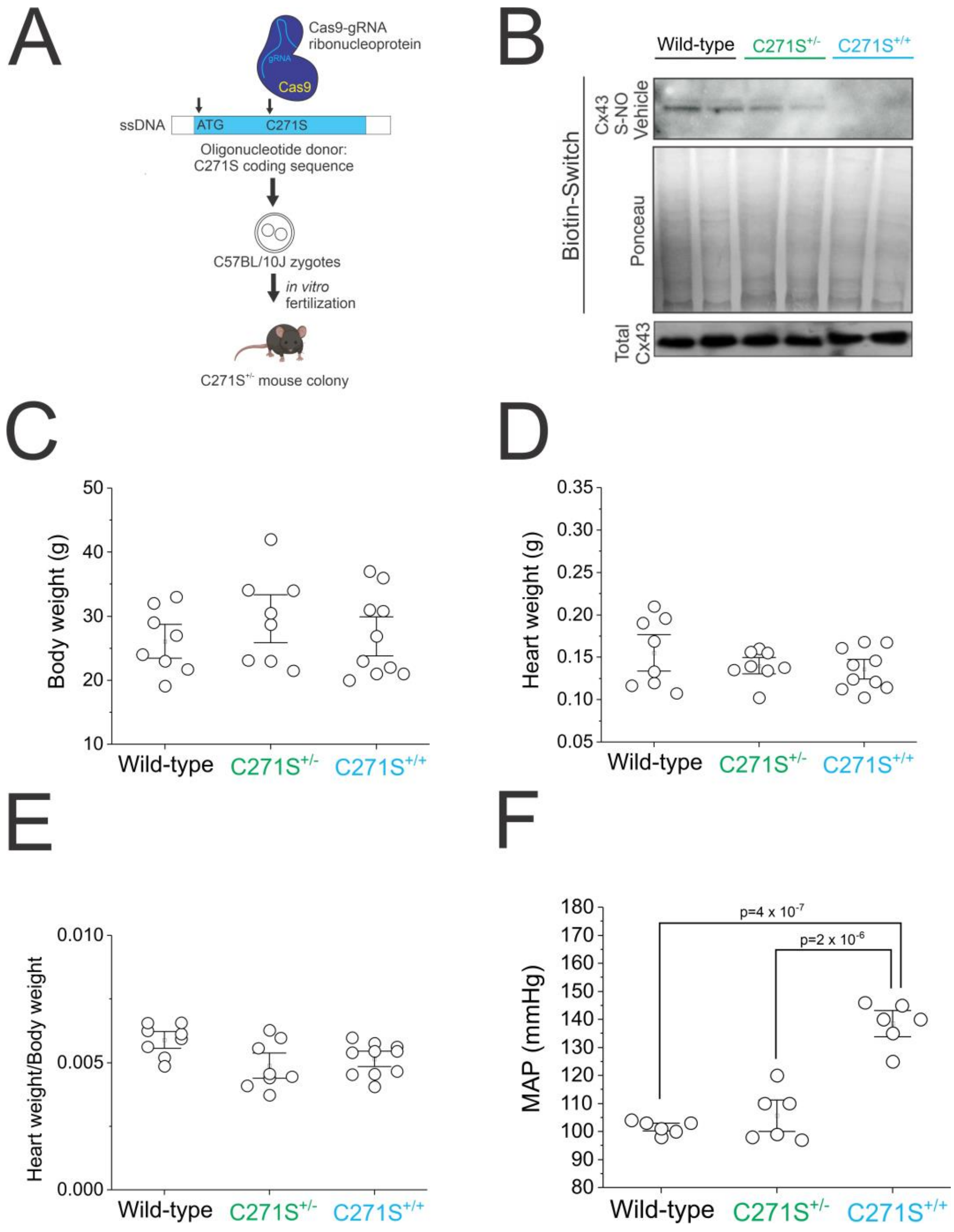


**Supplementary Figure 2. Characterization of Cx43$^{C271S}$ mutants.** (A) Schematic representation of the knock-in mouse model Cx43 developed using CRISPR-Cas9 genome editing, in which the site of S-nitrosylation of Cx43, cysteine 271 (C271), was replaced by a serine (C271S). (B) Representative western blots showing levels of S-nitrosylated Cx43 proteins (Cx43 S-NO) in heterozygous Cx43:C271S$^{+/-}$ (C271S$^{+/-}$) and homozygous Cx43:C271S$^{+/+}$ (C271S$^{+/+}$) mouse hearts. The middle blot is the Ponceau staining of the total pull-down for S-nitrosylated proteins. The lower blot shows total expression levels of Cx43 in the heart. Each lane represents an individual mouse heart. (C) Quantification of body weight, heart weight (D), and heart weight/body weight ratio (E) from wild-type (black), C271S$^{+/-}$ (green), and C271S$^{+/+}$ (blue) mice. (F) Quantification of mean arterial pressure (MAP) measured by a telemetry system in the carotid artery of mice. Each dot represents an independent animal. Group comparisons were made using a one-way ANOVA followed by Tukey’s post hoc test. Data are presented as means ± SEM.

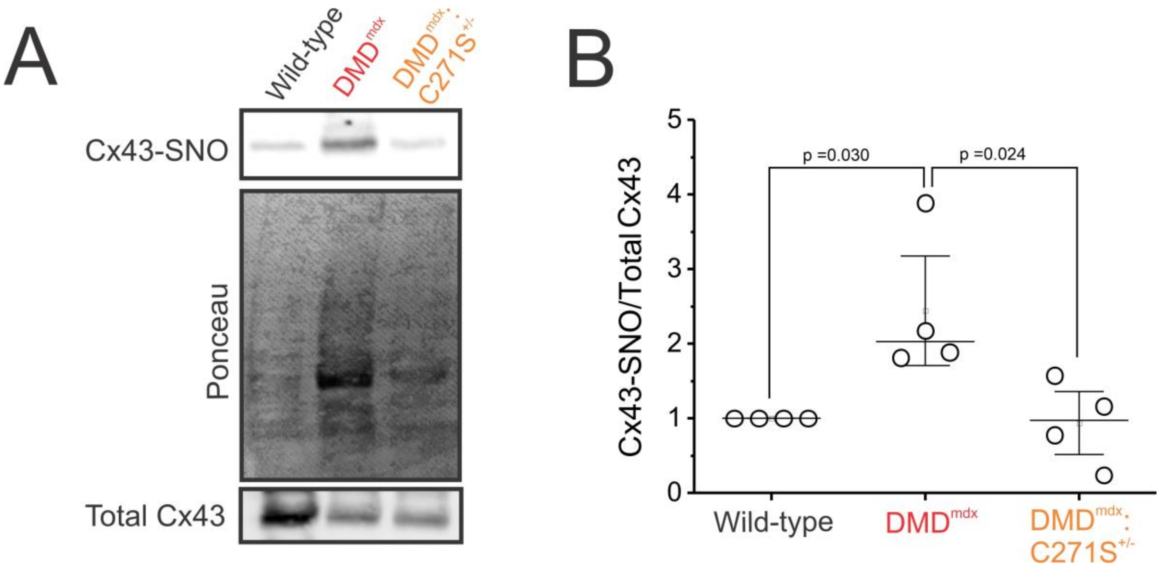


**Supplementary Figure 3. DMD$^{mdx}$:C271S$^{+/-}$ knockin mice exhibit reduced S-nitrosylation of cardiac Cx43 proteins.** (A) The top membrane was loaded with S-nitrosylated proteins isolated from heart samples from wild-type, DMD$^{mdx}$, and DMD$^{mdx}$:C271S$^{+/-}$ mice using the biotin switch assay. This membrane was probed for connexin 43 (Cx43). The middle panel shows the corresponding Ponceau staining, and the bottom panel depicts the total cardiac proteins blotted for Cx43. (B) Quantification of blots, presented as the ratio of S-nitrosylated Cx43 to total Cx43 protein. Each dot represents an independent animal. Group comparisons were made using a one-way ANOVA followed by Tukey’s post hoc test. Data are presented as means ± SEM.

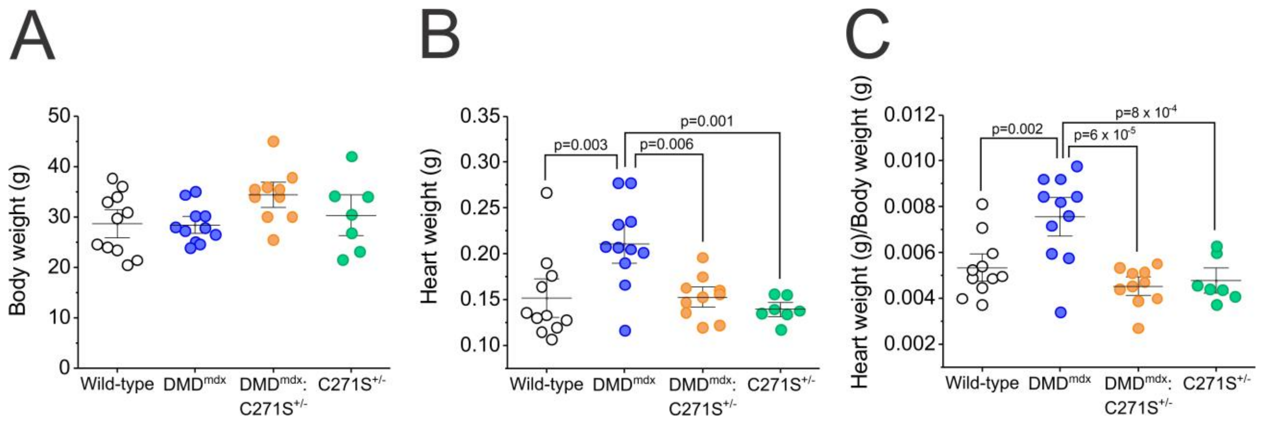


**Supplementary Figure 4. Cardiac hypertrophy observed in $DMD^{mdx}$ is prevented in $DMD^{mdx}$:$C271S^{+/}$ mice.** (A) Quantification of body weight, heart weight (B), and heart weight/body weight ratio (C) from wild-type (white circles), $DMD^{mdx}$ (blue circles), $DMD^{mdx}$:$C271S^{+/-}$ (orange circles), and $C271S^{+/-}$ (green circles) mice. Each dot represents an independent animal. Group comparisons were made using a one-way ANOVA followed by Tukey's post hoc test. Data are presented as means ± SEM.
